# Applying a conservation-based approach for predicting novel phosphorylation sites in eukaryotes and evaluating their functional relevance

**DOI:** 10.1101/2025.01.09.632054

**Authors:** Anton Kalyuzhnyy, Patrick A Eyers, Claire E Eyers, Eric W Deutsch, Andrew R Jones

## Abstract

Protein phosphorylation, a key post-translational modification, is central to cellular signalling and disease pathogenesis. The development of high-throughput proteomics pipelines has led to the discovery of large numbers of phosphorylated protein motifs and sites (phosphosites) across many eukaryotic species. However, the majority of phosphosites are reported from human cellular sources, with most species having only a few experimentally confirmed or computationally predicted phosphosites. Furthermore, only a small fraction of the characterised human phosphoproteome has an annotated functional role. A common method of predicting functionally relevant phosphosites is conservation analysis. However, extensive evolutionary studies involving large numbers of species are scarce. In this study, we explore the conservation of 20,751 confident human phosphosites across 100 eukaryotic species. We link the observed conservation patterns to the functional relevance of phosphosites and investigate the evolution of associated protein domains and kinases. We identify several protein functions that are likely regulated by phosphorylation, including ancient functions conserved across all life and relatively new functions only conserved in species closely related to humans. We demonstrate the importance of conservation analysis in identifying organisms suitable as biological models for studying conserved signalling pathways relevant to human biology and disease. Finally, we use human protein sequences as a reference to apply conservation analysis and predict over 1,000,000 potential phosphosites in analysed eukaryotes. Our results can ultimately be used to improve proteome annotations of several species and direct downstream research surrounding the evolution and functional relevance of eukaryotic phosphosites.

## Introduction

### The extent of protein phosphorylation in eukaryotes

In proteomics, kinase-regulated protein phosphorylation is perhaps the most important and frequently observed post-translational modification (PTM)^[1]^ which is well-studied in relation to cell signalling pathways and disease across all life^[2–4]^. The extent of protein phosphorylation in various eukaryotic species has been highlighted by several genome sequencing studies which revealed >500 kinase-encoding genes in humans^[5–8]^, >1,000 in *Arabidopsis thaliana*^[9]^, 240 in *Drosophila melanogaster*^[10]^ and 120 genes in *Saccharomyces cerevisiae*^[11]^. Furthermore, the development and optimisation of high-throughput proteomics pipelines such as tandem mass spectrometry (LC-MS/MS) has led to the discovery of large numbers of specific phosphorylated protein motifs and sites, focussing primarily on the phosphorylation of canonical (established) serine (Ser), threonine (Thr) and tyrosine (Tyr) residues^[12–17]^. Newly identified phosphorylation sites (phosphosites) are characterised and compiled in several publicly available resources, including PhosphoSitePlus (PSP)^[18]^, PeptideAtlas (PA)^[19]^, dbPAF^[20]^, The Plant PTM Viewer^[21]^ and PTMeXchange^[22–25]^, all of which accumulate post-translational modification data from eukaryotic species.

However, to our knowledge, most public databases do not account for phosphosite false discovery rate across large proteomics datasets, which results in the accumulation of false positive identifications^[23, 26]^. Therefore, the number of “*real*” phosphosites in those resources may be much lower than reported and researchers are advised to carefully evaluate evidence reporting phosphosites. Nevertheless, our previous in-depth analysis of phosphorylation data from commonly used resources such as PSP^[18]^ and PA^[19]^ revealed that the count of “*real*” phosphosites in humans alone may exceed 80,000^[23]^. We also established a set of over 20,000 “gold standard” phosphosites which had extensive positive identification evidence in both PSP and PA^[23]^. Phosphorylation remains an active area of research, with new phosphosites being reported and annotated as proteomics studies multiply.

### Predicting conservation and functional relevance of phosphosites

Despite numerous phosphosites being reported and extensive ongoing research surrounding the role of phosphorylation in proteomics, only a small fraction of the currently characterised human phosphoproteome has an annotated functional role^[18, 27]^. This is because the rate of phosphosite discovery is far greater than the rate at which each individual site or motif can be analysed experimentally. Furthermore, it has been proposed that a significant portion of phosphosites may be “*decoy*” sites or have no regulatory function at all^[28, 29]^. The difficulty in distinguishing functionally significant phosphorylated regions from those that do not contribute to protein function is exacerbated by the added complexity of proteins having multiple phosphorylated sites within their sequence, as well as several kinase enzymes being able to phosphorylate multiple sites^[30, 31]^.

A common and effective method for predicting functionally significant phosphorylated protein regions is site-conservation analysis. At its simplest, conservation analysis compares the amino acid sequence of a protein in question to the sequences of its homologues and identifies local regions of similarity which may have a common functional implication amongst the compared proteins^[32]^. Conservation analysis of genes and proteins often plays a central role in research surrounding model organisms and how they can be used to study human biology and disease^[33, 34]^. This is highlighted by various studies of organisms such as flies^[35]^, worms^[36]^, yeast^[37]^ and mammals^[36, 38]^ that uncovered novel molecular pathways and demonstrated a direct functional connection of those pathways to human biology by analysing conservation of the involved proteins^[34, 36]^.

It is generally hypothesised that functionally significant phosphosites would be highly conserved because their mutations to non-phosphosites would alter protein function and ultimately hinder evolutionary selection^[39, 40]^. Several studies demonstrated that Ser, Thr and Tyr phosphosites are indeed significantly more conserved compared to non-phosphosites in general^[27, 40, 41]^. Our previous profiling analysis of the human phosphoproteome also revealed that phosphosites are on average more conserved than sites with no phosphorylation evidence within the same protein sequences^[23]^. Multiple cross-species studies of phosphosites successfully characterised their conservation and linked it to functions such as cell cycle maintenance and metabolism^[42–44]^. However, extensive evolutionary studies which investigate phosphosite conservation across large numbers of species are scarce^[27, 45]^. This study provides a deeper insight into general evolutionary and functional trends surrounding human phosphosites by calculating and analysing phosphosite conservation within specific groups of eukaryotic species (vertebrates and invertebrates, mammals only, primates only, etc.) and understanding their functional relevance within those groups. We also analyse functional enrichment of human protein sets with different phosphosite conservation patterns by using clusterProfiler^[46]^ which is an open-source, user-friendly R package that offers comprehensive analysis and visualisation of enriched functions. Additional functional annotations are mapped using DAVID online tool^[47]^ which allows to extract functional terms from bioinformatics databases such as UniProtKB^[48]^, KEGG^[49]^, SMART^[50]^ and InterPro^[51]^. Furthermore, we establish conservation patterns of prevalent human protein domains by linking phosphosite conservation to the data from the database Pfam^[52]^ which offers accessible large-scale bulk downloads of accurate domain annotations within protein sequences. Finally, to further understand the evolution of phosphorylation, we analyse general conservation patterns of associated human Ser/Thr protein kinases.

### Predicting novel phosphosites in eukaryotes

The studies of phosphorylation sites and kinases are usually limited to a specific set of species, with the most frequent phosphosite discoveries being made in humans, followed by several model organisms such as mouse, flies, worms, yeasts and Arabidopsis^[18, 19]^. As a result, the number of reported phosphosites varies between species, with most eukaryotes having little to no evidence of either experimentally confirmed or computationally predicted phosphosites^[18, 45, 48]^. For example, in the January 2023 build of PhosphoSitePlus database, there were only 24 characterised eukaryotes with any reported phosphorylation data in low or high-throughput studies, where species such as ferret, starfish and buffalo only had a single known phosphosite^[18]^. The process of identifying phosphosites with LC-MS/MS and subsequent proteome annotation can be expensive and time-consuming, and its complexity also depends on the size of the proteome being annotated, as well as the availability of accurate experimental and genomic data for the species from which the proteome originates^[53, 54]^. As a result, phosphoproteomics studies are not attempted for the majority of the species. In fact, most experiments are designed and funded with the purpose of benefiting vertebrate (notably human) lives, as well as improving the understanding of human biology and diseases, which is often achieved by analysing a specific set of model organisms that have conserved functions in humans^[33, 34, 36]^.

Various computational tools such as ConSurf^[55]^, ACES^[56]^, Ensembl Compara^[57]^, NetPhos^[58]^ incorporate algorithms which analyse sequence conservation to predict functionally relevant sites within given protein sequences. Some phosphosite annotations in UniProtKB are also propagated from sequence similarity between a query sequence and a well-annotated homologous sequence with experimentally confirmed phosphosites, provided that the phosphorylated residue and the surrounding motif is conserved in the homologous sequence^[48]^. The propagations are usually limited between closely related species from the same taxonomic group and can be further validated if the kinase responsible for modifying the target is also conserved between the species^[48]^. In this study, we expand the scope of eukaryotic phosphosite predictions by mapping conserved “gold standard” phosphosites from the reference human proteome to aligned sites in potentially homologous sequences from 100 eukaryotic species.

## Methods

### Establishing phosphosite conservation patterns

To ensure that only the most confident phosphosites were utilised in this study, we exclusively focused on our previously characterised “gold standard” set of phosphosites (i.e., phosphosites which had at least 5 pieces of positive identification evidence from high or low throughput proteomics experiments referenced in PSP and PA databases)^[23]^. In total, the analysed set contained 16,978 Ser, 2,747 Thr and 986 Tyr sites across 5,709 proteins (Table S1). For those sites, we calculated conservation percentage scores across 100 eukaryotic species (Table S2) by applying the computational pipeline for evolutionary conservation analysis described in our previous study^[23]^. In brief, each human protein sequence with target phosphosites was used as a query in a BLASTp search (BLAST+ 2.10.0 version)^[59]^ against all 100 eukaryotic proteomes (complete proteomes in FASTA format are summarised in Supplementary file *proteomes.zip*). The BLAST output was processed to extract a top significant hit (E-value ≤1E-5) from each species for each human query. Human sequences were then aligned with their matched hits using MUSCLE (version 3.8.31)^[60]^. From each alignment, percentage conservation scores were calculated for every target human phosphosite and their +1 residue, taking into account any Ser/Thr substitutions in aligned sequences, whereby a site was counted if a Thr in its sequence was aligned with a Ser in the target human sequence and vice versa.

To establish phosphosite evolutionary patterns within eukaryotes, the data was processed to determine percentage conservation scores within specific groups of species such as primates (*n*=18), other mammals (*n*=32), birds (*n*=12), fish (*n*=5), reptiles (*n*=4), amphibians (*n*=2), insects/invertebrates (*n*=11), fungi (*n*=4), plants (*n*=7) and protists (*n*=5). In addition, conservation scores were calculated for broader groups such as animals (*n*=84), vertebrates (*n*=73) and mammals (*n*=50). To allow downstream protein-level functional analysis, the average conservation of all Ser/Thr and Tyr phosphosites across each described species group was calculated for each protein in the “gold standard” set.

Taxonomic relationships between the selected 100 eukaryotic species were displayed with a phylogenetic tree built with NCBI’s Taxonomy Browser tool^[61]^ using relevant UniProtKB proteome ID numbers as inputs (Table S2). The resulting phylogenetic tree was annotated and visualised using MEGA (version 10.2.2)^[62]^ and iTOL (version 5)^[63]^. Additional silhouette images within the resulting tree were obtained from PhyloPic database (https://www.phylopic.org/).

All proteins in the analysis (*n*=5,709) were then grouped into ten conservation clusters based on similarities in their Ser/Thr and Tyr phosphosite conservation patterns across the described species groups. The clustering was performed automatically using pheatmap package (version 1.0.12)^[64]^ in R programming software^[65]^, which applied the Euclidean distance method to assess similarity in phosphosite conservation patterns between target proteins and group them into specific clusters. The resulting protein clusters were presented as heatmaps and each cluster was manually named with an appropriate descriptive label which corresponded to the most observed conservation pattern within the cluster (i.e., at least 50% of proteins within the cluster had to match the label description in terms of their conservation patterns). The assigned labels characterised phosphosite conservation across species groups as “*High*” (phosphosites are ≥75% conserved within described species group) or “*Medium*” (phosphosites are ≥50% conserved within described species group). Similar clustering analysis was performed on individual sites, where target Ser/Thr and Tyr phosphosites were grouped according to their percentage conservation across the species groups. To highlight the diversity of conservation patterns identified for target phosphosites, multiple sequence alignments for randomly selected proteins that accurately matched the description of the corresponding clusters of interest and had a characterised function in UniProtKB were annotated and visualised in Jalview (version 2.11.2.3)^[66]^.

To compare the conservation of phosphosites located within ordered and disordered protein regions, the likelihood of target sites to be found within those regions was first predicted using metapredict v2.0^[67]^, which produced a score from 0 to 1, where a threshold of >0.3 was used for establishing disordered regions. The conservation of Ser, Thr and Tyr phosphosites was then compared between ordered and disordered protein regions using boxplots created using Tukey’s method^[68]^, where each box extended from the first quartile to the third quartile of the data and the whiskers extended from the box to the farthest data point lying within 1.5x the interquartile range from the box.

### Functional enrichment analysis of conservation clusters

Each protein cluster was analysed with R package clusterProfiler (version 4.4.1)^[46]^ to determine the functional enrichment of proteins with certain Ser/Thr and Tyr phosphosite conservation patterns against a control background of all analysed proteins (*n*=5,709) with at least one phosphosite from the “gold standard” set. For each cluster, a maximum of the 10 most enriched Gene Ontology (GO) terms were selected and visualised as dot plots, where the number of enriched proteins for a given GO term was provided and the statistical significance of the enrichment was measured with adjusted p-values. Clusters were counted as enriched for a particular GO term if their Benjamini–Hochberg adjusted p-value was <0.1. The functional enrichment analysis of the target protein clusters was extended by utilising DAVID online tool (version 6.8)^[47]^ and again using all analysed proteins (*n*=5,709) as a control background. For each cluster, the top 10 (where possible) significant (Benjamini–Hochberg adjusted p-value <0.1) functional terms with the highest percentage of proteins mapped were identified, with any near synonymous terms being filtered out. In addition to mapping target clusters to GO terms, DAVID analysis was also used to determine whether the clusters were enriched for any KEGG pathways^[49]^, UniProtKB keywords^[48]^ and annotations from domain databases such as SMART^[50]^ and InterPro^[51]^.

### Linking phosphosite conservation with protein domains

In order to link protein domains which encompass target Ser, Thr and Tyr phosphosites with their conservation patterns across the species groups, domain data was extracted from Pfam database (version 35.0; November 2021 release)^[52]^ and cross-referenced with the site-level conservation data by protein’s UniProtKB ID tag and site position within the protein sequence. In order to identify domains for which phosphosites with specific conservation patterns were enriched, fold enrichment (i.e., enrichment factor) was calculated for each domain against a control background of all phosphosites mapped to any Pfam domains. To calculate fold enrichment, a standard equation for enrichment analysis was applied as follows:

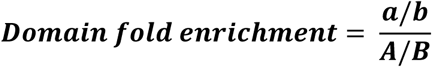

*a* = Count of phosphosites with a specific conservation pattern mapped to domain X
*b* = Count of all phosphosites with that specific conservation pattern
*A* = Count of all phosphosites mapped to domain X
*B* = Count of all phosphosites in the background distribution (i.e., all phosphosites mapped to any protein domain)

The domain data was then filtered to only include domains with at least two mapped phosphosites. The top 10 domains with the highest percentage of mapped sites from each Ser/Thr and Tyr conservation pattern were visualised and log_2_(fold enrichment) was presented for easier interpretation.

### Conservation analysis of associated protein kinases

All kinase-substrate mappings for our target Ser/Thr phosphosites were obtained from a published atlas of substrate specificities for the majority of the human kinome which computationally ranked kinases in terms of their relative likelihood of phosphorylating several reported human phosphosites^[30]^. For each of the analysed phosphosites in our set, we selected the top ranked kinase match from the atlas. Several kinases were selected per site if they had the same maximum likelihood score.

To investigate conservation patterns of the matched kinases, we first identified their established orthologue groups by searching UniProtKB^[48]^ and extracting group identification numbers for the OrthoDB database (v11)^[69]^ which provides high quality orthologue annotations. Each mapped OrthoDB group was then processed to extract its complete list of unique species containing protein orthologues of the target kinases. To ensure consistency in terms of species selection relative to the phosphosite conservation analysis, the list of species per OrthoDB group was cross-referenced with our selection of 100 eukaryotes (Table S2) to ultimately determine the extent of kinase conservation. The kinases were clustered according to their conservation patterns within the established groups of eukaryotes by applying a similar method described above for the analysis of phosphosite conservation. Finally, the conservation scores of target Ser/Thr phosphosites and their associated kinases in the selected 100 eukaryotic species were compared using linear regression.

### Predicting phosphosites across eukaryotes

Multiple sequence alignments between target human proteins and their potential homologues from the selected species were processed to identify all amino acids in the matched sequences which were aligned with target Ser, Thr and Tyr phosphosites in the human sequences. In addition, we analysed proximal amino acids adjacent to the aligned sites at the +1 position which are often involved in facilitating substrate recognition by the kinase enzymes^[13, 70, 71]^. For every species in each alignment, if both the amino acid that is aligned with the human phosphosite and its +1 adjacent site were conserved in the human sequence (considering Ser/Thr substitutions), then that amino acid was predicted to be a phosphosite. In order to validate the resulting phosphosite predictions, phosphorylation data was extracted for mouse and Arabidopsis from PSP^[18]^ and Plant PTM Viewer^[21]^ databases, respectively, to determine how many of the predicted phosphosites in those species had experimental phosphorylation evidence. Additional validation of our phosphosite predictions was done by assessing the likelihood of a certain site in mouse being aligned with a human phosphosite and also having strong phosphorylation evidence (>5 pieces of evidence in PSP). The resulting phosphosite predictions across 100 eukaryotic species were summarised into an easily accessible file which can be used to support the annotations of eukaryotic proteomes.

## Results and Discussion

### Evolutionary and functional analysis of human phosphorylation sites

In this study, we evaluated conservation of human phosphosites across 100 eukaryotic species ranging from primates and other vertebrates to plants and fungi (Table S2). By analysing average Ser/Thr and Tyr phosphosite conservation from each target human protein, we were able to split the phosphoproteins into independent clusters according to their similarity in phosphosite conservation patterns within specific groups of eukaryotes. Overall, our analysis successfully identified distinct phosphosite conservation patterns of human proteins (Table 1; Fig 1; Table S1). The diversity of the established conservation patterns was further highlighted by multiple sequence alignments of individual human protein examples (Fig 2). The differences in conservation patterns of human phosphosites were a likely result of their functional relevance within groups of species in which they are conserved^[27, 39, 40]^. To investigate this further, we performed a functional enrichment analysis of proteins with different phosphosite conservation patterns to identify conserved functions which are potentially regulated by those phosphosites. Indeed, we found that some phosphosites were involved in ancient protein functions relevant to all life forms such as regulation of cell cycle and metabolism, while others were likely contributing to relatively novel functions (brain and muscle development, motility and immune response) which were only conserved in specific species groups more closely related to humans (mammals or primates, for example) (Fig 3 and Fig S1). We linked the conservation patterns of mapped phosphosites with their associated domains to further understand their evolution and functional relevance in eukaryotes. By identifying frequently observed (enriched) domains per conservation cluster (Fig 4), we provided further evidence to support our results from the functional enrichment analysis.

**Table 1.** Independent clusters of human phosphoproteins split according to their similarity in average Ser/Thr and Tyr phosphosite conservation patterns across established species groups. High and medium conservation indicates that phosphosites are ≥75% and ≥50% conserved across the species in each cluster, respectively.

| Conservation pattern | Phosphosite | Protein count | Phosphosite count | Cluster label |
| --- | --- | --- | --- | --- |
| High in all vertebrates except birds | Ser/Thr | 45 | 80 | A |
| High in animals only | Ser/Thr | 384 | 820 | B |
| High or medium in all species | Ser/Thr | 197 | 379 | C |
| High in primates only | Ser/Thr | 603 | 1,650 | D |
| High in mammals only | Ser/Thr | 1,100 | 4,573 | E |
| Medium in all vertebrates except birds | Ser/Thr | 528 | 1,827 | F |
| High in mammals and medium in reptiles | Ser/Thr | 327 | 993 | G |
| High in all vertebrates except fish | Ser/Thr | 837 | 3,636 | H1 |
| High in vertebrates only | Ser/Thr | 1,222 | 5,041 | H2 |
| Medium in all vertebrates except amphibians | Ser/Thr | 274 | 726 | H3 |
| High in all vertebrates and medium in plants | Tyr | 21 | 24 | I |
| High in all species | Tyr | 89 | 102 | J |
| Medium in all species except fungi and protists | Tyr | 27 | 40 | K |
| High in vertebrates only | Tyr | 190 | 313 | L1 |
| High in animals only | Tyr | 103 | 160 | L2 |
| Medium in primates only | Tyr | 10 | 10 | M |
| High in mammals only | Tyr | 69 | 85 | N |
| Medium in all vertebrates except fish and amphibians | Tyr | 67 | 82 | O1 |
| High in all vertebrates except fish | Tyr | 53 | 61 | O2 |
| Medium in all vertebrates except birds | Tyr | 74 | 109 | P |

**Figure 1.**
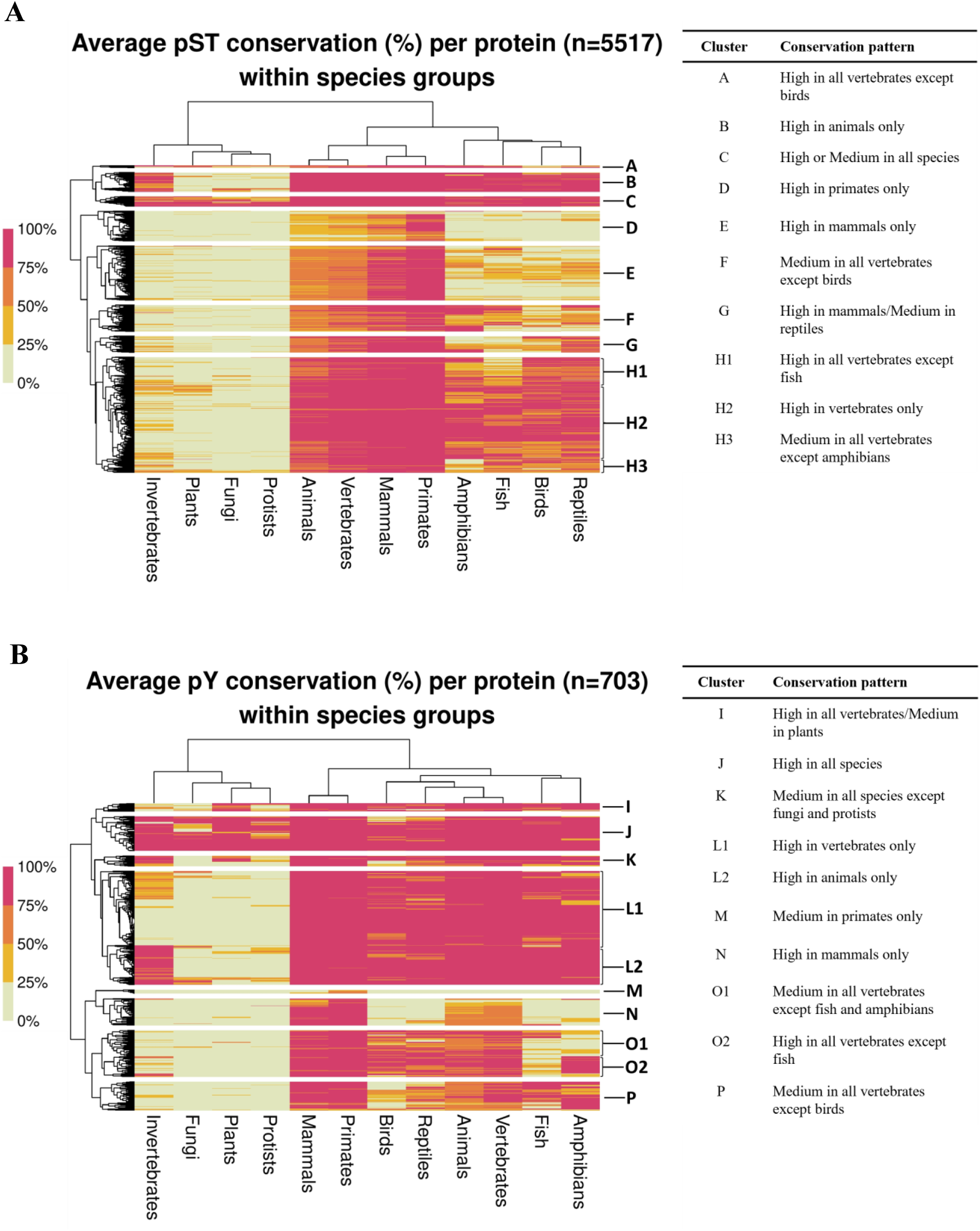
Conservation patterns of **(A)** Ser/Thr and **(B)** Tyr phosphosites from human proteins across the groups of eukaryotic species. Each row in the heatmap represents an individual human protein and its phosphosite conservation across specific species groups which are separated into columns. Conservation is scored as a percentage out of all species per group and reflected by a colour gradient divided at quarterly intervals. The proteins are clustered based on their similarity in conservation patterns using the Euclidean distance method. For each cluster, a label is assigned which describes the most observed conservation pattern (i.e., at least 50% of proteins in the cluster follow the described phosphosite conservation pattern), where high and medium conservation refers to conservation scores of ≥75% and ≥50%, respectively. The total number of analysed proteins containing target phosphosites is given by *n*.

**Figure 2.**
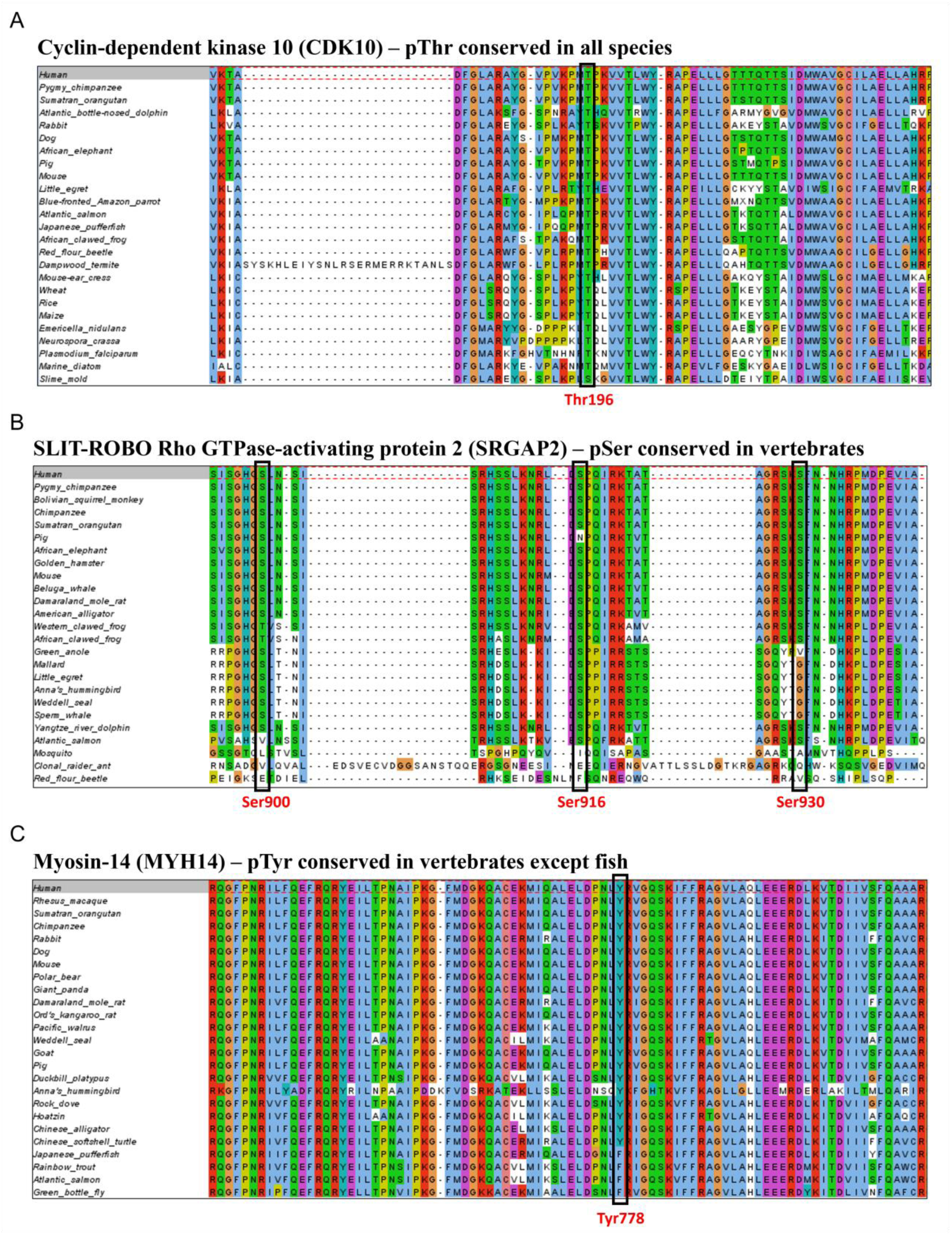
Sections of multiple sequence alignments of target human proteins with different conservation patterns of phosphosites across eukaryotic species. For each alignment, the sequence of the human protein containing the phosphosites is located at the top and the aligned protein sequences of potential orthologues from other species are given below. Phosphorylated sites are marked by black rectangle boxes and their location within the human sequence is provided underneath in red. Alignments were annotated in Jalview.

**Figure 3.**
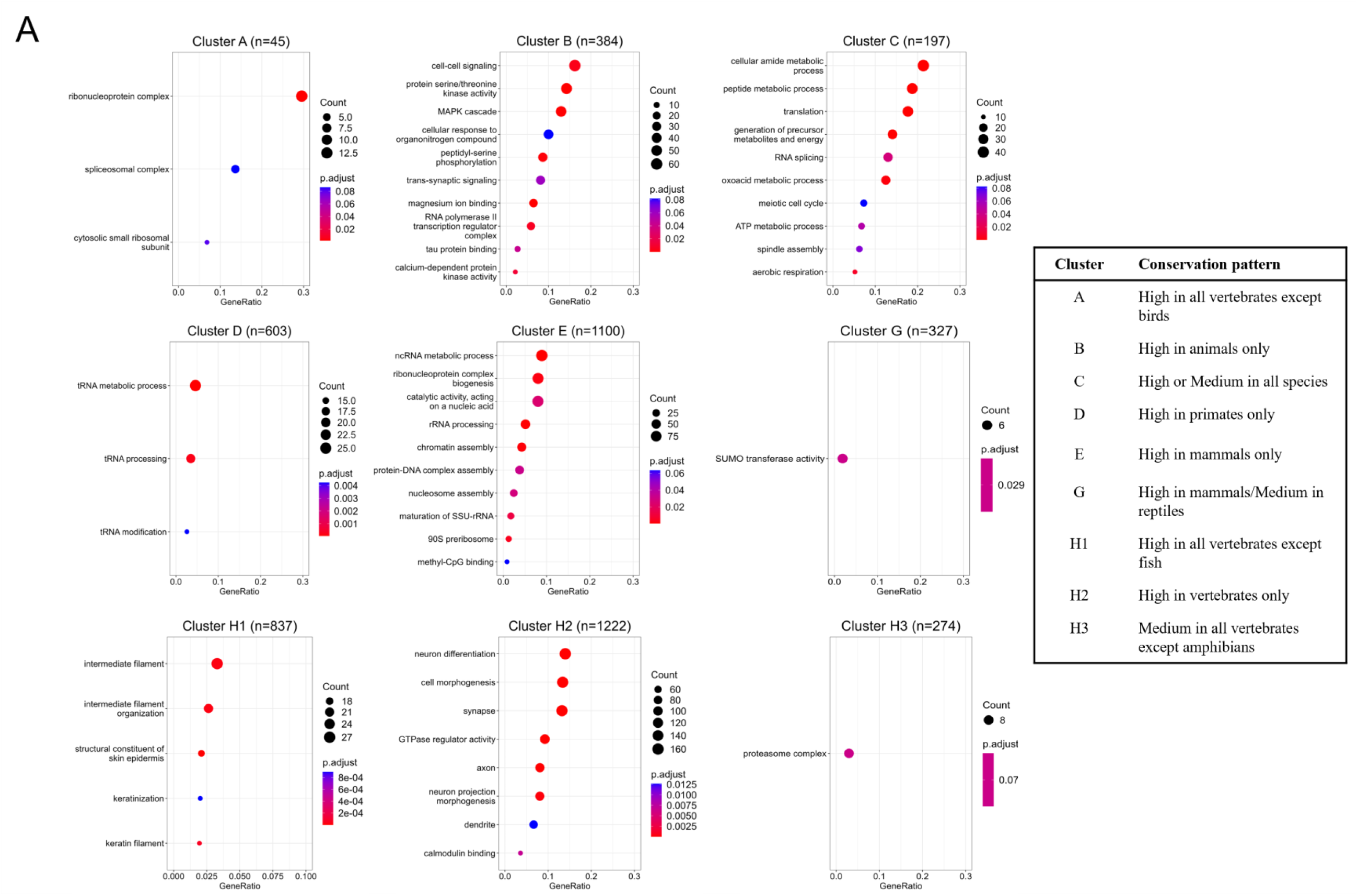
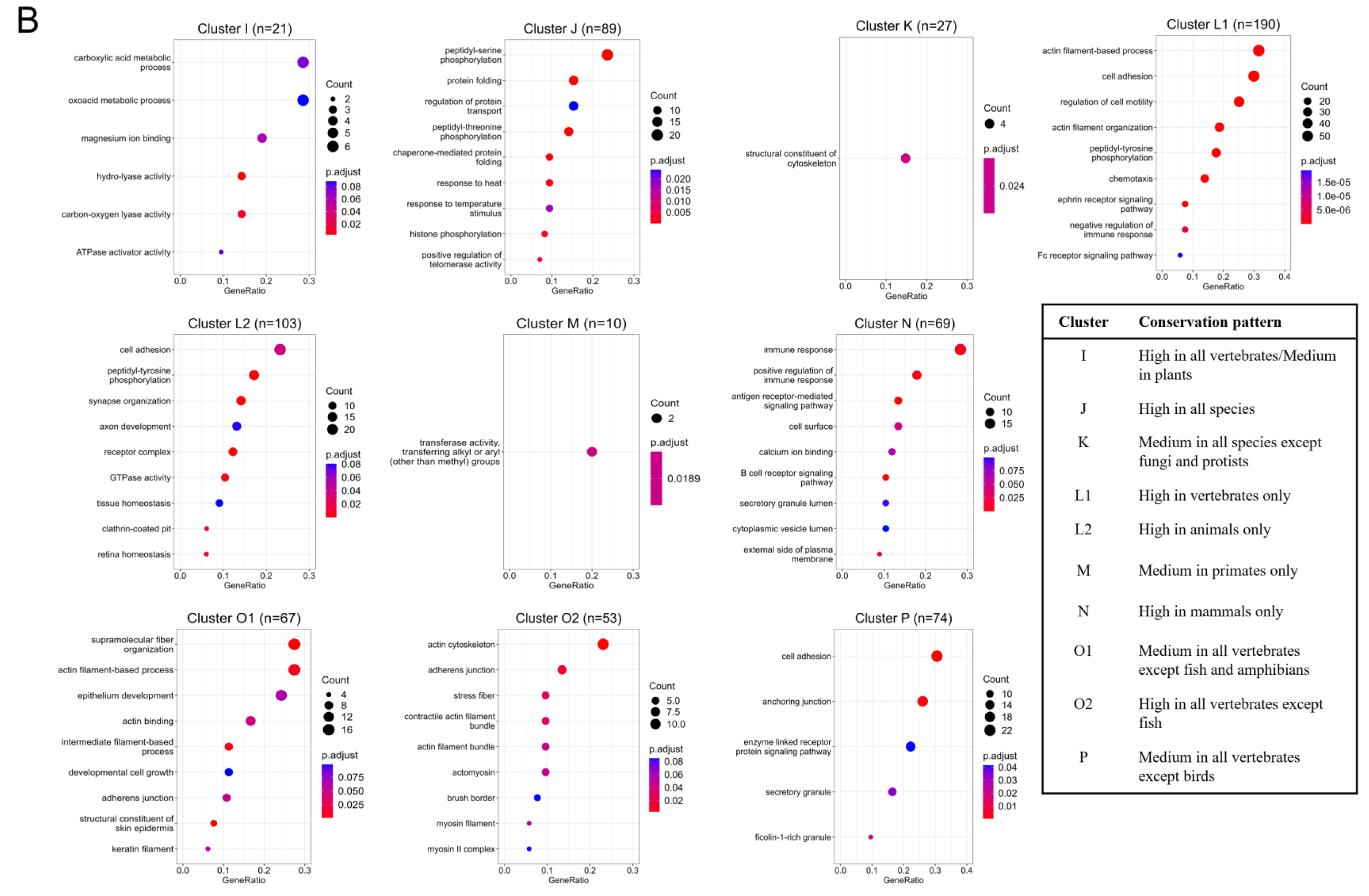
Functional enrichment for Gene Ontology terms of human phosphoproteins with different **(A)** Ser/Thr and **(B)** Tyr phosphosite conservation patterns. The enrichment is visualised with dot plots generated using clusterProfiler. In each dot plot, the dots represent protein sets enriched for a specific functional term described on the y-axis. The size of the dots reflects the number of proteins in the enriched set and the colour corresponds to the significance of the functional enrichment determined by Benjamini–Hochberg adjusted p-value. The position of the dots on the x-axis indicates the proportion of enriched proteins out of all analysed proteins in the protein set. The total number of proteins in each set is given by *n*.

**Figure 4.**
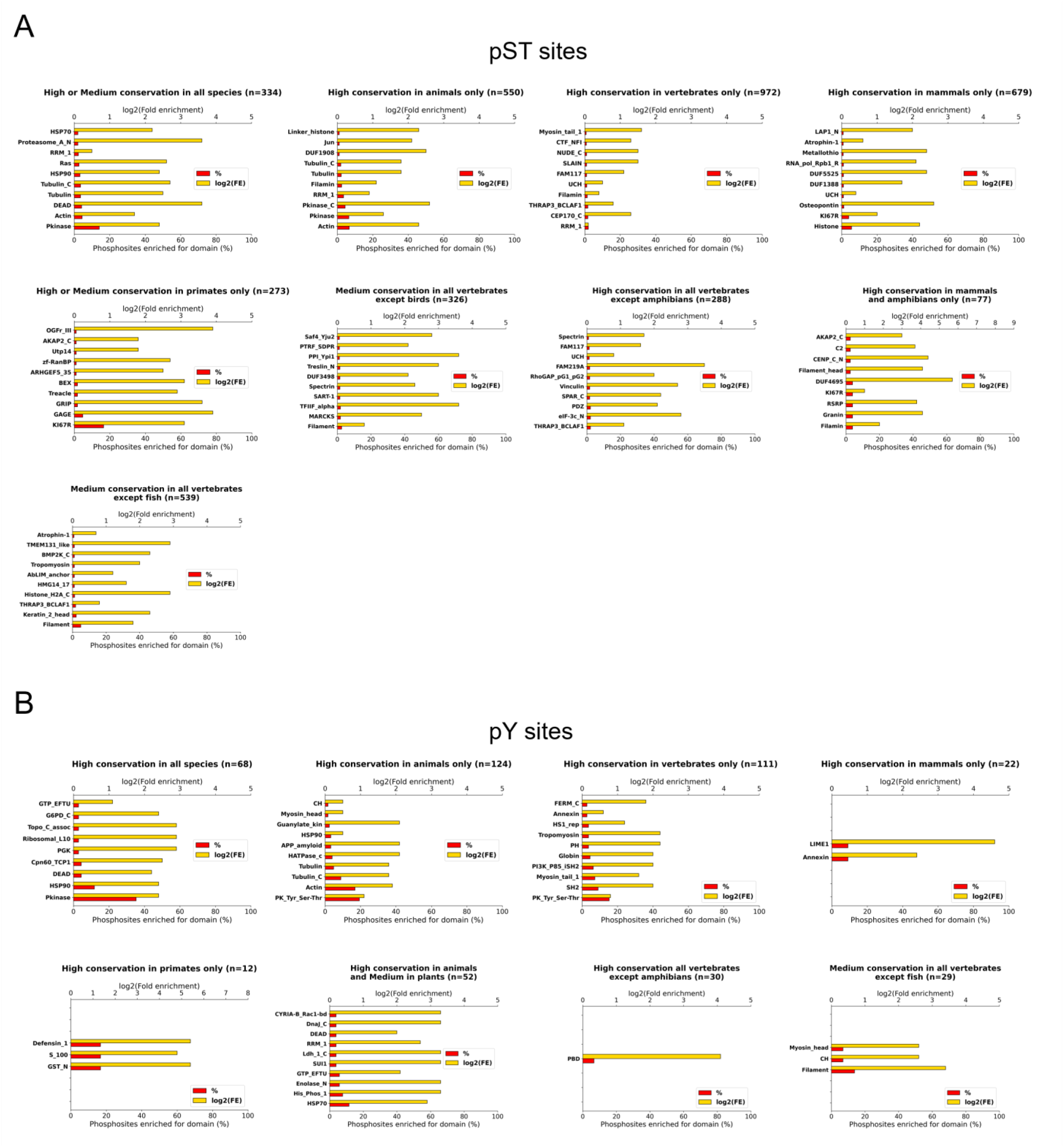
Protein domains from Pfam database (y-axis) for which **(A)** Ser/Thr and **(B)** Tyr phosphosites with specific conservation patterns across eukaryotes were enriched against a control background of all phosphosites mapped to any Pfam domains. For each conservation pattern, the percentage (%) of phosphosites with that pattern mapped to a specific domain is given, as well as the log2(fold enrichment). The total number of phosphosites with a particular conservation pattern mapped to any domain is presented by n.

Firstly, we identified 197 and 89 proteins in which Ser/Thr and Tyr phosphosites, respectively, were on average highly conserved in all species, ranging from mammals to plants and single-celled organisms (Fig 1A, cluster C; Fig 1B, cluster J). We linked those proteins to ancient molecular functions relevant across all life such as translation, metabolism, cell cycle regulation and heat response (Fig 3A, cluster C; Fig 3B, cluster J). For example, human Thr196 site in protein Q15131 (cyclin-dependent kinase 10; CDK10) was well-conserved across all species (Fig 2A) and its phosphorylation plays a role in promoting cell proliferation and transcription regulation^[72]^. This provides further evidence for the functional significance and conservation of CDK enzymes across life, with many CDK enzymes having been previously characterised across all species^[73]^. Additional DAVID analysis of proteins with phosphosites conserved in all species (Fig S1A, cluster C; Fig S1B, cluster J) revealed enrichment for acetylation which suggests that those post-translational modifications, along with phosphorylation, play a role in key biological functions across all life, potentially as part of crosstalk between PTMs^[74]^. Finally, phosphosites conserved across most species were linked to likely ancient protein domains found in several kinases, heat shock proteins, proteins associated with apoptosis (indicated by the enrichment of “*DEAD*” domains^[75]^), as well as actin and tubulin proteins (Fig 4). However, we do not rule out experimental biases meaning that different PTM sites might have been more heavily studied in conserved proteins involved in cell cycle and development.

In addition, we found numerous instances where human Ser/Thr phosphosites were conserved in broader species groups such as animals, vertebrates or mammals (Fig 1A, Table 1). For example, Ser900, Ser916 and Ser930 sites in protein O75044 (SLIT-ROBO Rho GTPase-activating protein 2; SRGAP2) were conserved in most vertebrates, but not in insects (Fig 2B). It is probable that the aligned insect sequences are not true orthologues of the human protein as indicated by the lack of any conserved motifs in the analysed phosphorylated region (Fig 2B). For human SRGAP2, there were also no significant matches found in BLAST (*E*-value ≤1E-5) from plant species, fungi and protists. This conservation pattern can be explained by the functional relevance of SRGAP2 in neuronal morphogenesis during the development of cerebral cortex necessary for complex brain functions, which are expectedly not present in insects, plants, fungi and protists^[76, 77]^. This connection was further highlighted by the functional enrichment analysis, which revealed a general enrichment of associated proteins for brain development-related terms such as “*neuron differentiation*”, “*synapse*” and “*axon*” (Fig 3A, cluster H2). Most importantly, the conservation of the identified pathways regulated by proteins with Ser/Thr phosphosites characterised in our analysis suggests that the selected vertebrates can be used as model organisms to study human brain development pathways.

Interestingly, we also found that 613 proteins (11%) in which phosphosites were only conserved in primates, indicating their potential relevance in relatively novel functions which diverged from other animals (Table 1; Fig 1A, cluster D; Fig 1B, cluster M). A functional enrichment analysis of those proteins inferred their involvement in tRNA processing and association with zinc finger-related terms including Kruppel-associated box (KRAB) zinc finger proteins (Fig 3A; Fig S1A, cluster D). In fact, previous studies also characterised groups of KRAB proteins which rapidly evolved in primates and adapted to regulate complex pathways involved in brain development^[78, 79]^. Furthermore, some proteins were characterised as G-antigen (GAGE) proteins (Fig S1A, cluster D). Those proteins have been known to play a regulatory role in primate germ cell development and were proposed as candidates for immunotherapy of cancer due to their expression in many cancer tissues^[80]^. The analysis of protein domains also linked Ser/Thr phosphosites conserved in primates to the “*GAGE*” domain (Fig 4B). Additionally, we found an enrichment of Ser/Thr phosphosites conserved in mammals for the “*KI67R*” domain which is known to be associated with genome stability and mammalian immune response against highly proliferative cells^[81]^. This domain was further enriched in primates suggesting its potential involvement in additional, primate-exclusive signalling pathways (Fig 4A). A similar pattern was found for Tyr phosphosites conserved in mammals or primates and enriched for protein domains known to be involved in immune response such as “*LIME1*”^[82, 83]^, “*Annexin*”^[84]^ and “*Defensin_1*”^[85]^, further emphasising that phosphorylation plays a key role in immune system pathways of higher eukaryotes (Fig 4B). The results highlight that identifying human phosphosites which are conserved in closely related species can extend the availability of genetically similar species for studying human neuronal and immune systems, as well as developing preclinical models for drug development.

Another noteworthy conservation pattern was found for 890 (16%) proteins in which phosphosites were conserved in most vertebrates except fish (Fig 1A, cluster H1; Fig 1B, cluster O2). The functional analysis of those proteins revealed strong enrichment for terms associated with keratin and myosin (Fig 3A and Fig S1A, cluster H1; Fig 3B and Fig S1B, cluster O2). Moreover, the domain analysis revealed enrichment for the “*Filament*” domain (Fig 4) involved in the formation of intermediate filaments^[86]^. Firstly, this suggests that the conserved phosphosites may be involved in the formation of intermediate filaments during keratinisation, a process required for the development of epidermis^[87]^. It is likely that those signalling pathways are absent in fish because they are not necessary for the survival in aquatic environment. This conservation pattern was further highlighted by evolutionary studies which similarly identified groups of keratin proteins only conserved in tetrapods (i.e., four-legged vertebrates which evolved later than fish) and linked those proteins to biological pathways involved in the development of tissues and organs such as skin, hair and nails necessary for protection against the friction caused by terrestrial movement^[88, 89]^. Secondly, the enrichment for myosin indicates the involvement of the associated proteins and their phosphosites in cell signalling and motility pathways likely absent in fish. One such protein is Q7Z406 (Myosin-14; MYH14), in which Tyr778 site was conserved across most vertebrates, but was mutated to phenylalanine in fish (Fig 2C). This evolutionary pattern was also confirmed in another study which identified several myosin domains that emerged as a result of divergent evolution between fish and tetrapods^[90]^. The example highlights the functional relevance of tyrosine phosphorylation in tetrapods and helps understand the evolution of established myosin-related functions such as cell signalling and motility^[91, 92]^. In addition, MYH14 has been previously linked to hearing loss in humans^[93]^ and the species in which the functional Tyr phosphosite was conserved can potentially be used as potential clinical models to further study its involvement in human disease.

We were unable to link some of our clusters to specific biological functions that might have helped explain the observed conservation patterns (Fig 3). This is likely a result of a small sample size of proteins involved in those clusters or weaker enrichment which did not yield any conclusive results. It is also possible that there were sequencing or annotation errors in proteomes of certain species used in our analysis or within the specific protein sequences aligned with our human targets. This could have in turn led to errors in resulting multiple sequence alignments, causing consequent inaccuracies in the characterisation of phosphosite conservation patterns and functional enrichment predictions as demonstrated by several studies^[94–96]^. Nevertheless, we were able to showcase the relevance of conservation analysis in inferring a range of different protein functions likely regulated by phosphorylation and selecting species which can be used as genetic models to support human studies.

### Linking phosphosite conservation to protein domains and regions

Having established specific conservation patterns of human phosphosites, we were able to identify relevant protein domains in which those phosphosites were located by using data from Pfam^[52]^. In total, we successfully mapped 4,576 out of 20,711 (22%) of our analysed Ser, Thr and Tyr phosphosites to 1,419 different protein domains and protein families characterised in the Pfam database (Table S1). It is likely that the unmapped phosphosites had an independent functional role outside of any given established protein domain or belonged to a recently discovered protein which has not yet been annotated in Pfam. Indeed, we found that the vast majority (88%) of the target phosphosites were found in disordered protein regions (Table S1; Fig S2), which aligns with previous structural evaluations investigating phosphosite localisation^[97–99]^ and can be explained by the need of functional phosphosites to readily regulate protein function through surface-based mechanisms.

We also found that on average, phosphosites involved in ordered protein regions were more conserved compared to those found in disordered regions (Fig S2) which is likely due to their involvement in phosphorylation hotspots within eukaryotic protein families that regulate ancient functions^[27]^. Furthermore, in our analysis we identified several protein domains which were likely involved in regulating a range of those ancient protein functions and contained phosphosites that were highly conserved across all or the majority of analysed species (Fig 4).

Interestingly, we found several phosphosites within domains of unknown function (DUFs) (Fig 4A) which are described in Pfam as families of uncharacterised proteins that do not currently have a known function^[100]^. This suggests that phosphorylation could play an important role within the associated DUFs and our results can ultimately be used to improve their functional annotation.

### Conservation of human kinases in kinomes

Out of 19,765 Ser and Thr sites from the “gold standard” set, we mapped 18,752 (95%) sites to 301 kinase candidates from the kinome atlas^[30]^ (Table S1). Out of the mapped sites, 543 (3%) sites had multiple kinase candidates with the same maximum match probability score in the atlas (Table S1). To discover the evolutionary relationships of the associated kinases, we linked them to the data in the OrthoDB database and identified 101 unique orthologue groups (Table S3). On average, in the mapped OrthoDB orthologue groups there were 2,964 orthologues (range: 97-8,739 proteins) from 709 different species (range: 83-1,238 species) (Table S3). To ensure consistency with phosphosite data analysis in terms of species selection and conservation pattern clustering, we only investigated kinase conservation out of the 100 analysed eukaryotes, whereby a kinase was considered conserved in the selected species if it was found in the OrthoDB group mapped to that kinase. In total, we found that 92 of our selected 100 species were also present in the OrthoDB analysis (Table S1), which allowed us to perform an analogous conservation clustering analysis for the kinases. We assumed that the kinases were not conserved in the missing 8 species or had significantly divergent sequences such as those found in some pseudokinases and atypical kinases that require structural-based informatics to define^[101–103]^.

In the clustering analysis, we found that the conservation signal was initially weak for birds, likely because the specific bird species selected in our analysis were not extensively studied in terms of kinase activity or had poor proteome annotations. Having excluded birds from the analysis, we obtained much clearer and more interpretable kinase conservation results (Fig S3). Similarly, the conservation signal was generally weak for the protist group, which was likely because two out of five analysed protists, *D. discoideum* and *T. pseudonana* were not found in any of the mapped OrthoDB groups (Table S1). Nevertheless, we identified several interesting conservation patterns for the target human kinases, including those conserved across all species and those found exclusively in vertebrates or in the broader group of higher eukaryotes (Fig S3).

For example, several members of the human G protein-coupled receptor kinase (GRK) family had orthologues in other animals, suggesting their involvement in signal transduction pathways associated with animal evolution (Fig S3). Previous evolutionary analysis of GRK enzymes concluded that they likely evolved in animals to allow adequate rapid signalling responses^[104]^. In addition, we found multiple human kinases which were conserved across most eukaryotic species, indicating their likely involvement in regulating ancient protein functions through phosphorylation (Fig S3). For example, members of CDK, p21-activated kinase (PAKs) and MAPK families regulate cell cycle and other proliferative pathways known to be conserved across eukaryotic life^[105–107]^. The results highlight the general diversity of kinase conservation across eukaryotes.

To further explore the evolution of protein phosphorylation, we attempted to compare the conservation patterns between kinases and their associated phosphosites, but did not find any strong associations (Fig S4). For example, when analysing kinases which were conserved in most of our analysed eukaryotes (Fig S3, cluster C4), we found that several of their associated phosphosites were also strongly conserved. Conversely, we found that there were many human phosphosites which had weak conservation (for example, only conserved in primates) but were phosphorylated by ancient kinases such as cyclin-dependent kinase (CDK) or mitogen activated protein kinase (MAPK) (Table S1). This suggests that kinases can regulate ancient proteins involved in essential life functions and those that have evolved more recently through specific molecular mechanisms in higher eukaryotes.

It is possible that the lack of strong signal between phosphosites and their kinases could be due to how the kinases were mapped and our analysis also initially depended on the accuracy of the kinase mappings in the kinome atlas^[30]^. For this study, we selected the top scoring kinase hits from the kinome atlas for our target phosphosites. However, it is also possible that multiple kinases can phosphorylate the same phosphosite and conversely, a single kinase can phosphorylate multiple sites. In addition, our methods of estimating orthologous relationships for kinases and their target phosphoproteins were fundamentally different. As such, any individual associations (or lack thereof) between the conservation patterns of phosphosites and their mapped kinases based on the current data may not be definitive and may not fully consider the complexity of the biological signal surrounding phosphorylation that is yet to be fully understood. Nevertheless, we provide a comprehensive starting point for any follow-up investigations of the co-evolution between kinases and their protein substrates.

### Predicting phosphorylation sites in eukaryotes

We used high-quality human phosphosites (Table S1) as a reference set to infer phosphorylation in other eukaryotes by studying multiple sequence alignments between target human protein sequences and the aligned sequences from selected eukaryotes. In total, we predicted >830,000 Ser, >140,000 Thr and >56,000 Tyr potential phosphosites in the analysed species (Fig 5, Table S4). The majority of human phosphosites can be propagated to primates, likely due to their closest evolutionary relationship with humans, leading to similarities in protein sequences and common functional relevance. As expected, with increasing evolutionary distance from humans, the proportion of mapped sites generally decreased (Fig 5). In addition, we were able to predict hundreds of potential phosphosites in lower eukaryotes such as plants, fungi and protists based on their sequence alignment with human sites and likely involvement in common functions regulated by phosphorylation (Fig 5, Table S4).

**Figure 5.**
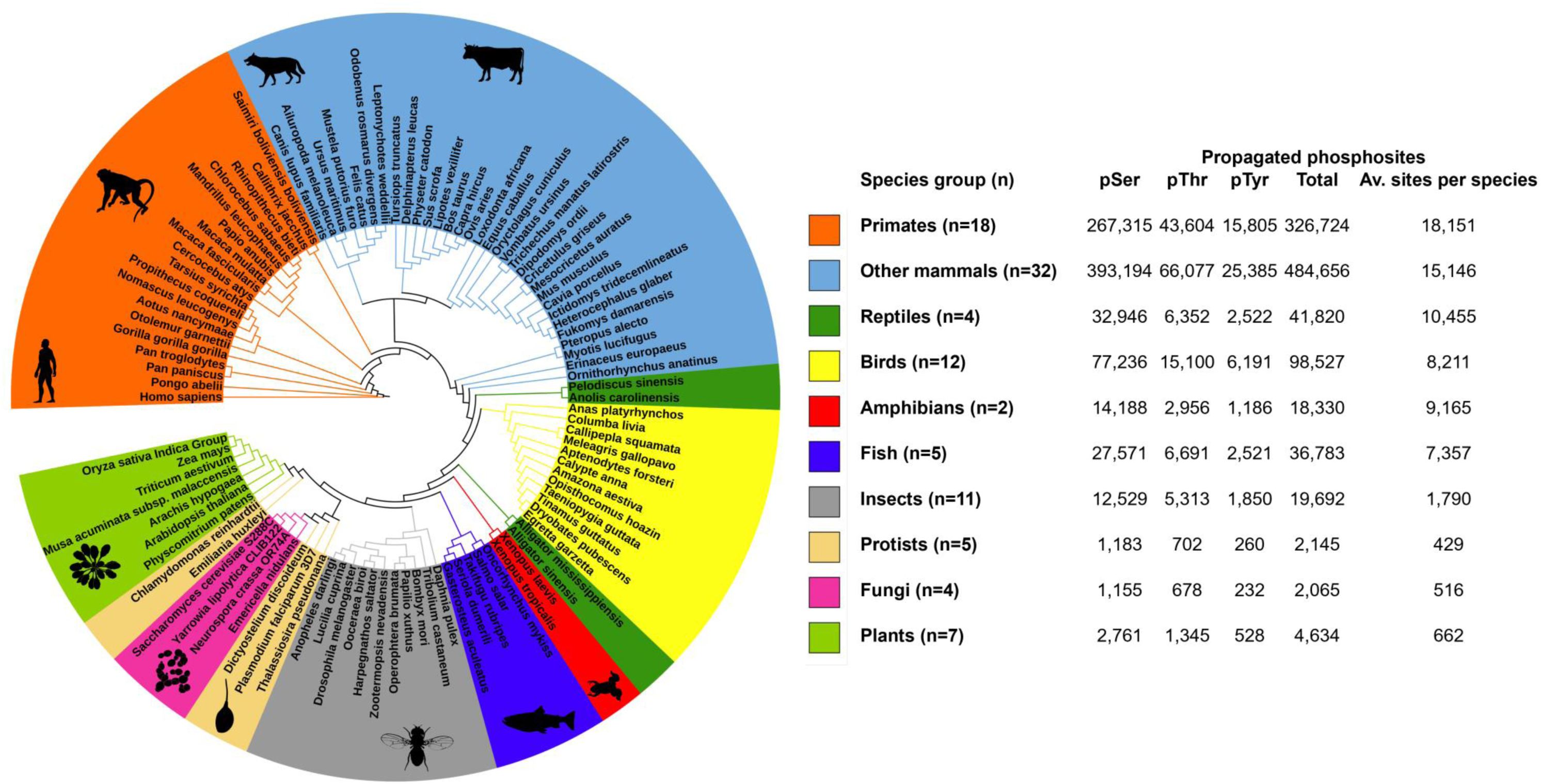
The phylogenetic relationship between the groups of eukaryotic species involved in the conservation analysis of human proteins and the resulting counts of propagated phosphosites from humans to other eukaryotes.

To validate our phosphosite predictions, we investigated how many of our predicted sites in species such as mouse and Arabidopsis had any actual experimental identification evidence in phosphorylation databases PSP^[18]^ and Plant PTM Viewer^[21]^, respectively. We found that 82% and 75% of our predicted Ser/Thr and Tyr phosphosites in mouse had reported experimental evidence in PSP, respectively (Table S4). Out of those sites, 61% had at least 5 pieces of phosphorylation evidence (Table S4), indicating a confident set of phosphosites with a low false discovery rate. By taking into account the total number of Ser, Thr and Tyr phosphosites in the mouse proteome and the overall number of phosphosites reported in PSP for mouse, we estimated enrichment factors of 14 and 43 for identifying confident phosphorylation sites that had any experimental evidence in PSP and at least 5 pieces of evidence, respectively, against a probability of identifying those sites by random chance. In other words, our method is much more likely to identify confident phosphosites in mouse than if they were selected at random. Furthermore, we found that 35% of our predicted Ser, Thr and Tyr phosphosites in Arabidopsis were supported by experimental evidence in Plant PTM Viewer resource (Table S4). The proportion of predicted Arabidopsis phosphosites mapped to experimental evidence was lower than for mouse predictions, likely because many of the target Arabidopsis protein sequences were missing phosphosite annotations or because many sequences have not yet been analysed experimentally and reported in Plant PTM Viewer. Nevertheless, our results suggest that analysing multiple sequence alignments between human proteins with confident phosphosites and their top sequence hits in BLAST from other species can successfully predict confident phosphosites in those species and act as a starting point for further analysis.

## Conclusion

A major strength of our approach lies in the use of established high quality human phosphorylation data to explore the functional importance of phosphosites. In particular, this allowed us to infer the relevance of phosphosites in several ancient and novel protein functions, explore their specific roles and analyse their associated protein domains. Our results emphasise the importance of conservation analysis in predicting functional significance of phosphosites and identifying organisms that can be used as additional models to study conserved signalling pathways relevant to human biology and disease.

To further understand the evolution of phosphorylation across eukaryotes, we analysed the conservation of human kinase enzymes which were most likely responsible for phosphorylating our “gold standard” phosphosites. As with phosphosites, we found various kinase conservation patterns across our selected eukaryotes. However, we could not find any significant correlation between the general conservation of phosphosites and their top matched kinases, likely due to the complexities of underlying biological signals not considered in the current study. Nevertheless, our results provide a good starting point for individual evolutionary investigations of target phosphosites and their respective kinases, particularly involving the inference of signalling mechanisms between different eukaryotic species.

Finally, by using the “gold standard” human phosphosite set as reference, we predicted over 1,000,000 potential Ser, Thr and Tyr phosphosites across various eukaryotic species ranging from primates and other mammals to plants, fungi and protists. We envisage that researchers would use our “gold standard” human phosphosites (Table S1) and the predicted phosphosites in other eukaryotes (Table S4) as a starting point to ultimately improve protein sequence annotations in different eukaryotic species and direct further research involving the use of those species as biological models for studying conserved cell signalling pathways in which phosphorylation plays an important role in health and disease. The development of deep-learning tools such as AlphaFold3^[108]^ for predicting protein structures in the context of phosphorylation makes structural informatics one obvious setting for this to occur.

