## Supplementary_Material for "Applying a conservation-based approach for predicting novel phosphorylation sites in eukaryotes and evaluating their functional relevance": Table S2. Proteomes summary.docx

| Organism name from Uniprot | Organism ID | Proteome ID | Gene count |
| --- | --- | --- | --- |
| *Homo sapiens* (Human) | 9606 | UP000005640 | 21152 |
| *Pan troglodytes* (Chimpanzee) | 9598 | UP000002277 | 23006 |
| *Pan paniscus* (Pygmy chimpanzee) (Bonobo) | 9597 | UP000240080 | 21221 |
| *Gorilla gorilla gorilla* (Western lowland gorilla) | 9595 | UP000001519 | 21795 |
| *Pongo abelii* (Sumatran orangutan) (Pongo pygmaeus abelii) | 9601 | UP000001595 | 21999 |
| *Nomascus leucogenys* (Northern white-cheeked gibbon) (Hylobates leucogenys) | 61853 | UP000001073 | 20762 |
| *Macaca mulatta* (Rhesus macaque) (Strain: 17573) | 9544 | UP000006718 | 21211 |
| *Cercocebus atys* (Sooty mangabey) (Cercocebus torquatus atys) | 9531 | UP000233060 | 20883 |
| *Rhinopithecus bieti* (Black snub-nosed monkey) (Pygathrix bieti) | 61621 | UP000233180 | 20855 |
| *Chlorocebus sabaeus* (Green monkey) (Cercopithecus sabaeus) | 60711 | UP000029965 | 19141 |
| *Macaca fascicularis* (Crab-eating macaque) (Cynomolgus monkey) (Strain: CE-4) | 9541 | UP000009130 | 17395 |
| *Papio anubis* (Olive baboon) | 9555 | UP000028761 | 21568 |
| *Saimiri boliviensis boliviensis* (Bolivian squirrel monkey) | 39432 | UP000233220 | 19365 |
| *Callithrix jacchus* (White-tufted-ear marmoset) | 9483 | UP000008225 | 19679 |
| *Aotus nancymaae* (Ma's night monkey) | 37293 | UP000233020 | 20372 |
| *Tarsius syrichta* (Philippine tarsier) | 1868482 | UP000189704 | 19956 |
| *Otolemur garnettii* (Small-eared galago) (Garnett's greater bushbaby) | 30611 | UP000005225 | 19451 |
| *Propithecus coquereli* (Coquerel's sifaka) (Propithecus verreauxi coquereli) | 379532 | UP000233160 | 17882 |
| *Ictidomys tridecemlineatus* (Thirteen-lined ground squirrel) (Spermophilus tridecemlineatus) | 43179 | UP000005215 | 18446 |
| *Cavia porcellus* (Guinea pig) (Strain: 2N) | 10141 | UP000005447 | 18253 |
| *Mus musculus* (Mouse) (Strain: C57BL/6J) | 10090 | UP000000589 | 22287 |
| *Oryctolagus cuniculus* (Rabbit) (Strain: Thorbecke inbred) | 9986 | UP000001811 | 19904 |
| *Cricetulus griseus* (Chinese hamster) (Cricetulus barabensis griseus) (Strain: CHO K1 cell line) | 10029 | UP000001075 | 23882 |
| *Fukomys damarensis* (Damaraland mole rat) (Cryptomys damarensis) | 885580 | UP000028990 | 20405 |
| *Mesocricetus auratus* (Golden hamster) | 10036 | UP000189706 | 20468 |
| *Dipodomys ordii* (Ord's kangaroo rat) | 10020 | UP000081671 | 19728 |
| *Heterocephalus glaber* (Naked mole rat) | 10181 | UP000006813 | 21445 |
| *Myotis lucifugus* (Little brown bat) | 59463 | UP000001074 | 19749 |
| *Canis lupus familiaris* (Dog) (Canis familiaris) (Strain: Boxer) | 9615 | UP000002254 | 20271 |
| *Ovis aries* (Sheep) (Strain: Texel) | 9940 | UP000002356 | 21218 |
| *Sus scrofa* (Pig) (Strain: Duroc) | 9823 | UP000008227 | 23228 |
| *Felis catus* (Cat) (Felis silvestris catus) (Strain: Abyssinian) | 9685 | UP000011712 | 19515 |
| *Ailuropoda melanoleuca* (Giant panda) | 9646 | UP000008912 | 19388 |
| *Pteropus alecto* (Black flying fox) | 9402 | UP000010552 | 19520 |
| *Erinaceus europaeus* (Western European hedgehog) | 9365 | UP000079721 | 19242 |
| *Equus caballus* (Horse) (Strain: Thoroughbred) | 9796 | UP000002281 | 21411 |
| *Bos taurus* (Bovine) (Strain: Hereford) | 9913 | UP000009136 | 23523 |
| *Mustela putorius furo* (European domestic ferret) (Mustela furo) (Strain: ID#1420) | 9669 | UP000000715 | 19909 |
| *Lipotes vexillifer* (Yangtze river dolphin) | 118797 | UP000265300 | 18846 |
| *Leptonychotes weddellii* (Weddell seal) | 9713 | UP000245341 | 20476 |
| *Ursus maritimus* (Polar bear) (Thalarctos maritimus) | 29073 | UP000261680 | 19366 |
| *Delphinapterus leucas* (Beluga whale) | 9749 | UP000248483 | 18472 |
| *Odobenus rosmarus divergens* (Pacific walrus) | 9708 | UP000245340 | 19319 |
| *Physeter macrocephalus* (Sperm whale) (Physeter catodon) | 9755 | UP000248484 | 18971 |
| *Tursiops truncatus* (Atlantic bottle-nosed dolphin) (Delphinus truncatus) | 9739 | UP000245320 | 17075 |
| *Loxodonta africana* (African elephant) (Strain: Isolate ISIS603380) | 9785 | UP000007646 | 20110 |
| *Trichechus manatus latirostris* (Florida manatee) | 127582 | UP000248480 | 19066 |
| *Ornithorhynchus anatinus* (Duckbill platypus) (Strain: Glennie) | 9258 | UP000002279 | 21678 |
| *Meleagris gallopavo* (Wild turkey) | 9103 | UP000001645 | 14171 |
| *Taeniopygia guttata* (Zebra finch) (Poephila guttata) | 59729 | UP000007754 | 17434 |
| *Anas platyrhynchos* (Mallard) (Anas boschas) | 8839 | UP000016666 | 15659 |
| *Manacus vitellinus* (golden-collared manakin) | 328815 | UP000053258 | 12707 |
| *Dryobates pubescens* (Downy woodpecker) (Picoides pubescens) | 118200 | UP000053875 | 13097 |
| *Tinamus guttatus* (White-throated tinamou) | 94827 | UP000053641 | 13377 |
| *Amazona aestiva* (Blue-fronted Amazon parrot) | 12930 | UP000051836 | 16092 |
| *Calypte anna* (Anna's hummingbird) (Archilochus anna) | 9244 | UP000054308 | 13267 |
| *Cuculus canorus* (common cuckoo) | 55661 | UP000053760 | 13582 |
| *Columba livia* (Rock dove) | 8932 | UP000053872 | 14726 |
| *Callipepla squamata* (Scaled quail) (Strain: Texas) | 9009 | UP000198323 | 16973 |
| *Aptenodytes forsteri* (Emperor penguin) | 9233 | UP000053286 | 13704 |
| *Opisthocomus hoazin* (Hoatzin) (Phasianus hoazin) | 30419 | UP000053605 | 12773 |
| *Lonchura striata domestica* (Bengalese finch) | 299123 | UP000197619 | 15248 |
| *Egretta garzetta* (Little egret) | 188379 | UP000053119 | 13489 |
| *Alligator mississippiensis* (American alligator) | 8496 | UP000050525 | 24656 |
| *Alligator sinensis* (Chinese alligator) | 38654 | UP000189705 | 19111 |
| *Anolis carolinensis* (Green anole) (American chameleon) (Strain: JBL SC #1) | 28377 | UP000001646 | 18550 |
| *Pelodiscus sinensis* (Chinese softshell turtle) (Trionyx sinensis) | 13735 | UP000007267 | 18116 |
| *Salmo salar* (Atlantic salmon) (Strain: double haploid) | 8030 | UP000087266 | 47721 |
| *Oncorhynchus mykiss* (Rainbow trout) (Salmo gairdneri) | 8022 | UP000193380 | 46447 |
| *Gasterosteus aculeatus* (Three-spined stickleback) | 69293 | UP000007635 | 20809 |
| *Tetraodon nigroviridis* (Spotted green pufferfish) (Chelonodon nigroviridis) | 99883 | UP000007303 | 19573 |
| *Takifugu rubripes* (Japanese pufferfish) (Fugu rubripes) | 31033 | UP000005226 | 20593 |
| *Xenopus laevis* (African clawed frog) (Strain: J) | 8355 | UP000186698 | 41590 |
| *Xenopus tropicalis* (Western clawed frog) (Silurana tropicalis) (Strain: Nigerian) | 8364 | UP000008143 | 24386 |
| *Daphnia pulex* (Water flea) | 6669 | UP000000305 | 30119 |
| *Tribolium castaneum* (Red flour beetle) (Strain: Georgia GA2) | 7070 | UP000007266 | 16579 |
| *Bombyx mori* (Silk moth) (Strain: p50T) | 7091 | UP000005204 | 14773 |
| *Anopheles stephensi* (Indo-Pakistan malaria mosquito) (Strain: Indian) | 30069 | UP000076408 | 11758 |
| *Anopheles darlingi* (Mosquito) | 43151 | UP000000673 | 10447 |
| *Drosophila melanogaster* (Fruit fly) (Strain: Berkeley) | 7227 | UP000000803 | 13788 |
| *Harpegnathos saltator* (Jerdon's jumping ant) (Strain: R22 G/1) | 610380 | UP000008237 | 15029 |
| *Ooceraea biroi* (Clonal raider ant) (Cerapachys biroi) | 2015173 | UP000053097 | 16497 |
| *Papilio xuthus* (Asian swallowtail butterfly) | 66420 | UP000053268 | 15265 |
| *Zootermopsis nevadensis* (Dampwood termite) | 136037 | UP000027135 | 14539 |
| *Operophtera brumata* (winter moth) | 104452 | UP000037510 | 16814 |
| *Lucilia cuprina* (Green bottle fly) (Australian sheep blowfly) (Strain: LS) | 7375 | UP000037069 | 14353 |
| *Saccharomyces cerevisiae* (strain ATCC 204508 / S288c) (Baker's yeast) | 559292 | UP000002311 | 6049 |
| *Emericella nidulans* (strain FGSC A4 / ATCC 38163 / CBS 112.46 / NRRL 194 / M139) (Aspergillus nidulans) | 227321 | UP000000560 | 10557 |
| *Arachis hypogaea (*Peanut*)* | 3818 | UP000289738 | 71122 |
| *Neurospora crassa* (strain ATCC 24698 / 74-OR23-1A / CBS 708.71 / DSM 1257 / FGSC 987) | 367110 | UP000001805 | 9759 |
| *Yarrowia lipolytica* (strain CLIB 122 / E 150) (Yeast) (Candida lipolytica) | 284591 | UP000001300 | 6449 |
| *Arabidopsis thaliana* (Mouse-ear cress) (Strain: cv. Columbia) | 3702 | UP000006548 | 27476 |
| *Oryza sativa subsp. indica* (Rice) (Strain: cv. 93-11) | 39946 | UP000007015 | 37344 |
| *Zea mays* (Maize) (Strain: cv. B73) | 4577 | UP000007305 | 39448 |
| *Triticum aestivum* (Wheat) (Strain: cv. Chinese Spring) | 4565 | UP000019116 | 130673 |
| *Physcomitrella patens subsp. patens* (Moss) (Strain: cv. Gransden 2004) | 3218 | UP000006727 | 30792 |
| *Emiliania huxleyi* (Pontosphaera huxleyi) (Strain: CCMP1516) | 2903 | UP000013827 | 35676 |
| *Dictyostelium discoideum* (Slime mold) (Strain: AX4) | 44689 | UP000002195 | 12739 |
| *Chlamydomonas reinhardtii* (Chlamydomonas smithii) (Strain: CC-503) | 3055 | UP000006906 | 17614 |
| *Thalassiosira pseudonana* (Marine diatom) (Cyclotella nana) (Strain: CCMP1335) | 35128 | UP000001449 | 11717 |
| *Plasmodium falciparum* (isolate 3D7) | 36329 | UP000001450 | 5448 |
