## Supplementary figures and images for "Applying a conservation-based approach for predicting novel phosphorylation sites in eukaryotes and evaluating their functional relevance"

### Figure S1. Functional Enrichment DAVID.png

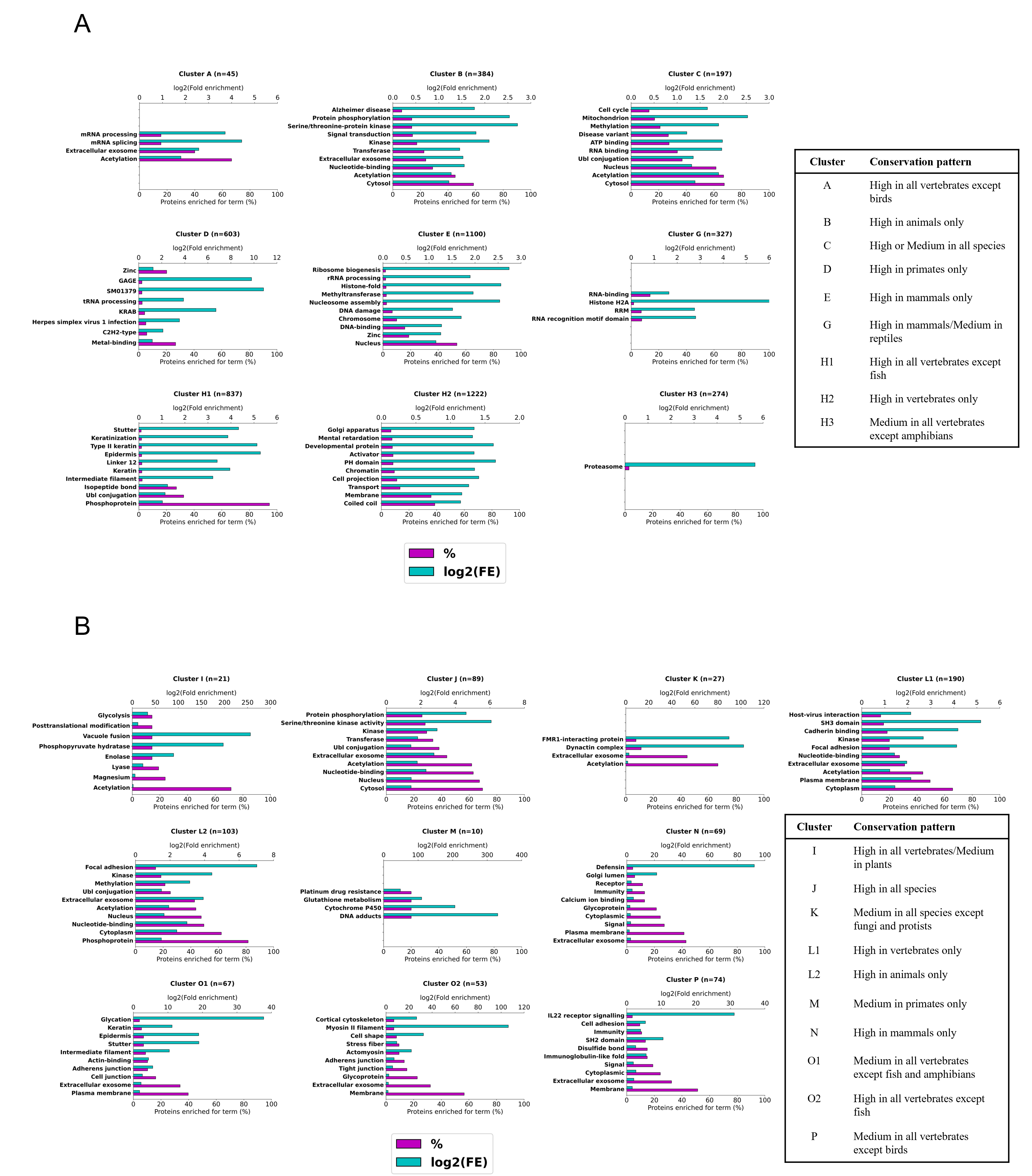

### Figure S2. Conservation vs protein regions.png

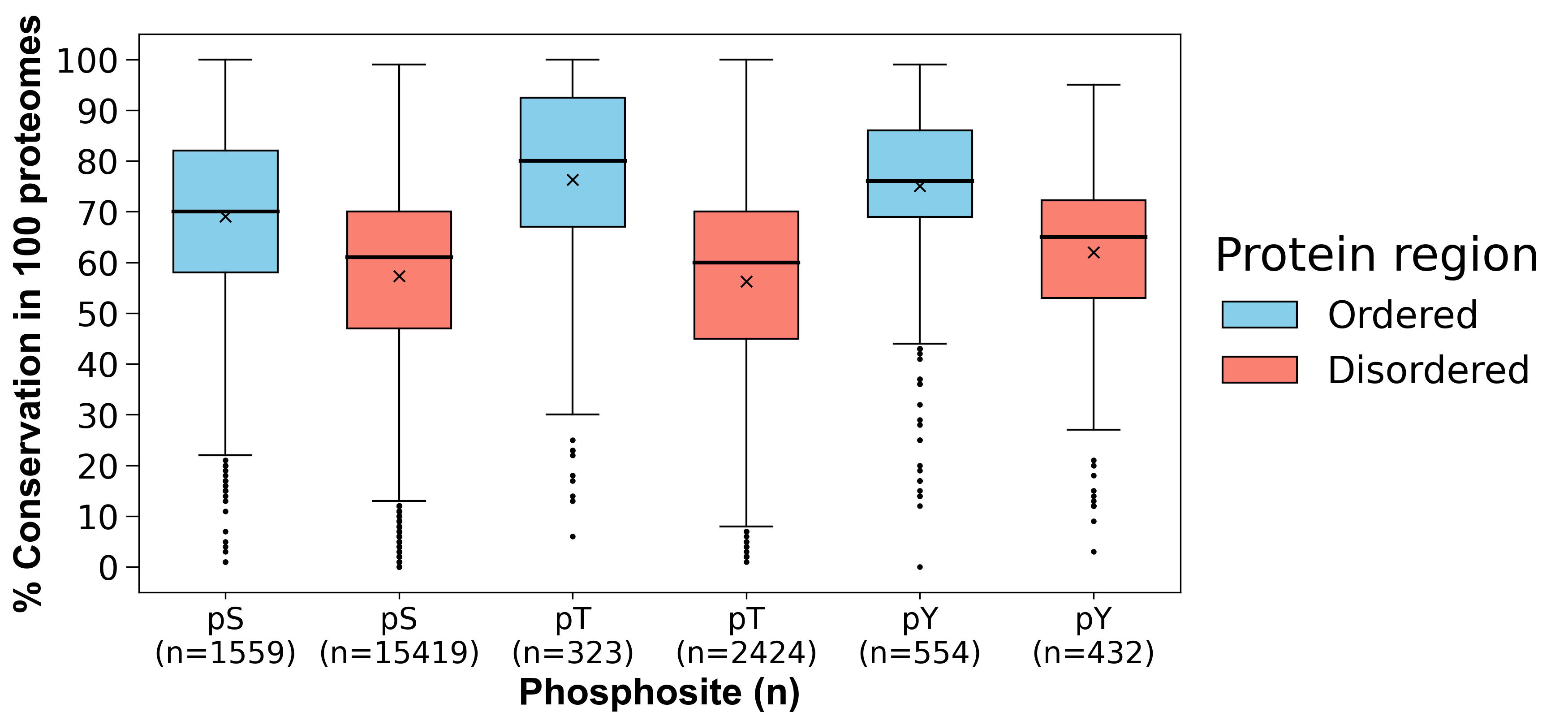

### Figure S3. Kinase conservation clusters.tif

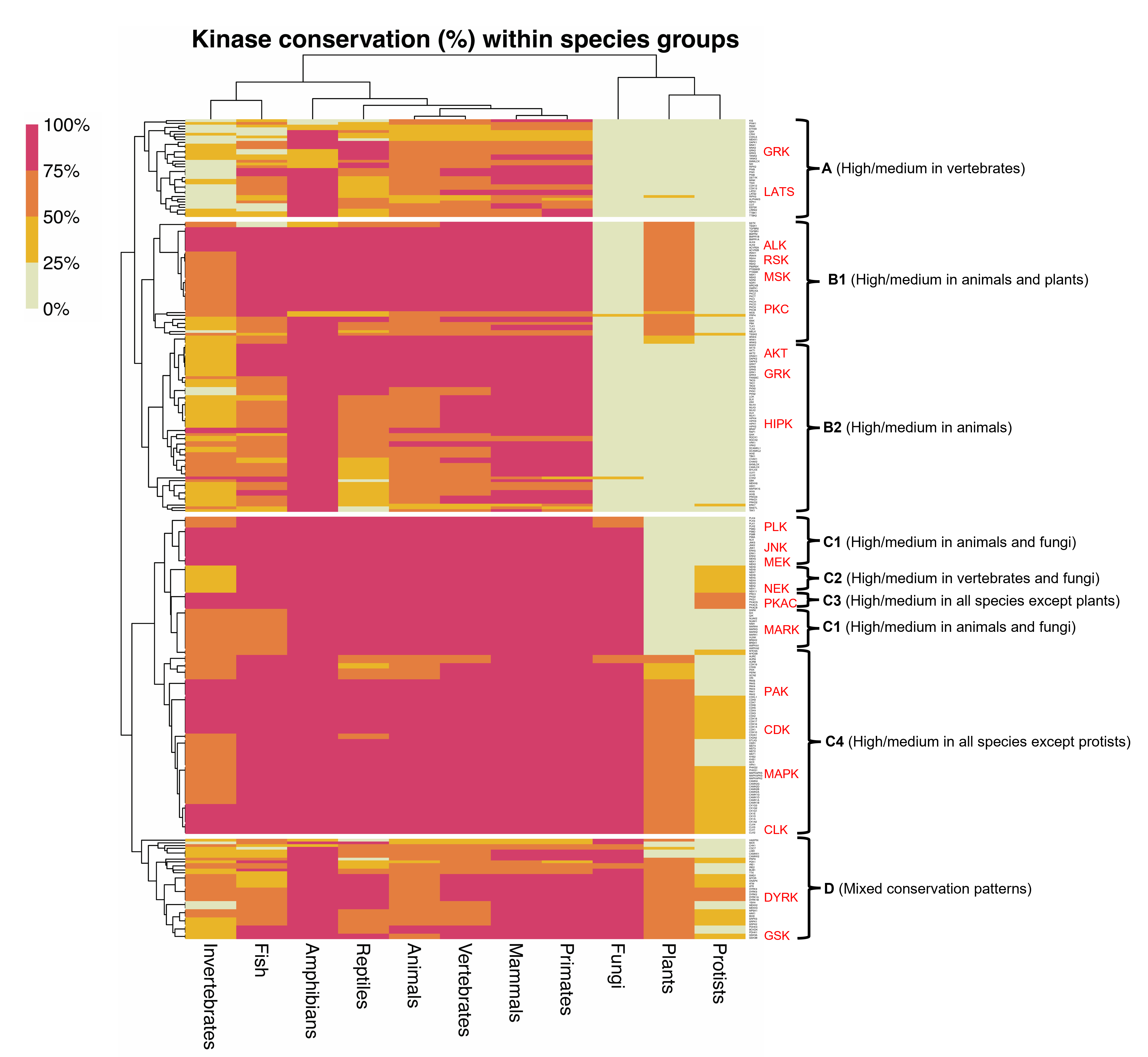

### Figure S4. Phosphosite Vs Kinase Conservation.png

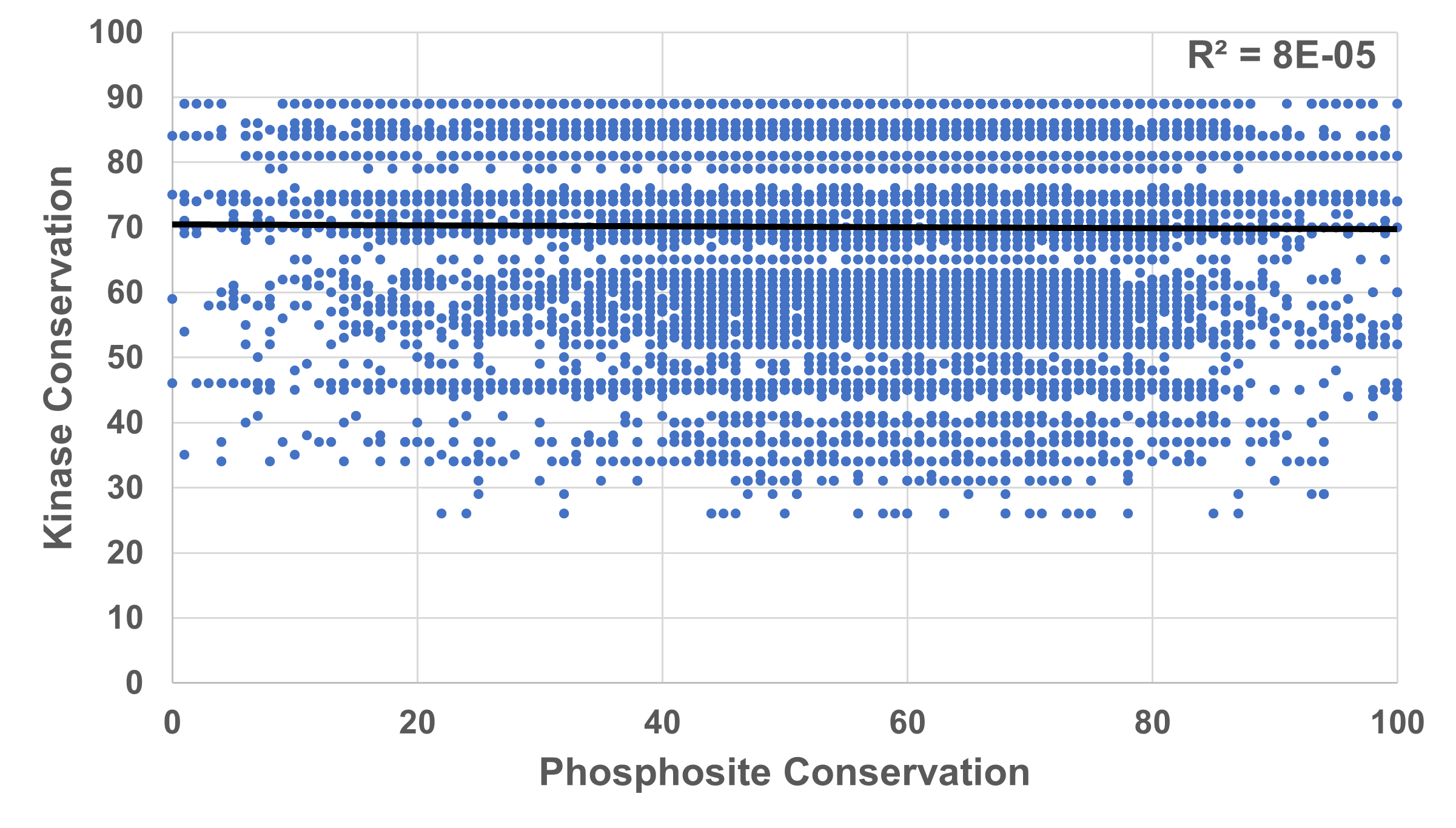
